# Opto-controlled C9orf72 poly-PR forms anisotropic condensates causative of TDP-43 pathology in the nucleus

**DOI:** 10.1101/2024.03.05.581933

**Authors:** Rachel E Hodgson, Jessica Rayment, Wan-Ping Huang, Anna Sanchez Avila, Tatyana A Shelkovnikova

## Abstract

Proteinaceous inclusions formed by *C9orf72* derived dipeptide-repeat (DPR) proteins are a histopathological hallmark in ~50% of familial amyotrophic lateral sclerosis/frontotemporal dementia (ALS/FTD) cases. However DPR aggregation/inclusion formation could not be efficiently recapitulated in cell models for four out of five DPRs. In this study, using optogenetics, we achieved chemical-free poly-PR condensation/aggregation in cultured cells, with spatial and temporal control. Strikingly, nuclear poly-PR condensates had anisotropic, hollow-centre appearance, resembling anisosomes formed by aberrant TDP-43 species, and their growth was limited by RNA. These condensates induced abnormal TDP-43 granulation in the nucleus without the activation of stress response. Cytoplasmic poly-PR aggregates that formed under prolonged light stimulation were more persistent than its nuclear condensates, selectively sequestered TDP-43 in a demixed state and surrounded spontaneous stress granules. Our data suggest that poly-PR anisotropic condensation in the nucleus, causative of nuclear TDP-43 dysfunction, may constitute an early pathological event in C9-ALS/FTD. Anisosome-type condensates may represent a more common cellular pathology in neurodegeneration than previously thought.

**Highlights:**

- Optogenetics can be used to model *C9orf72* DPR condensation in cultured cells.
- Opto-PR forms hollow nuclear condensates, and RNA limits their growth by fusion.
- Opto-PR condensation leads to stress-independent TDP-43 pathology in the nucleus.
- Cytoplasmic poly-PR assemblies are persistent and selectively sequester TDP-43.

**Graphical abstract:** 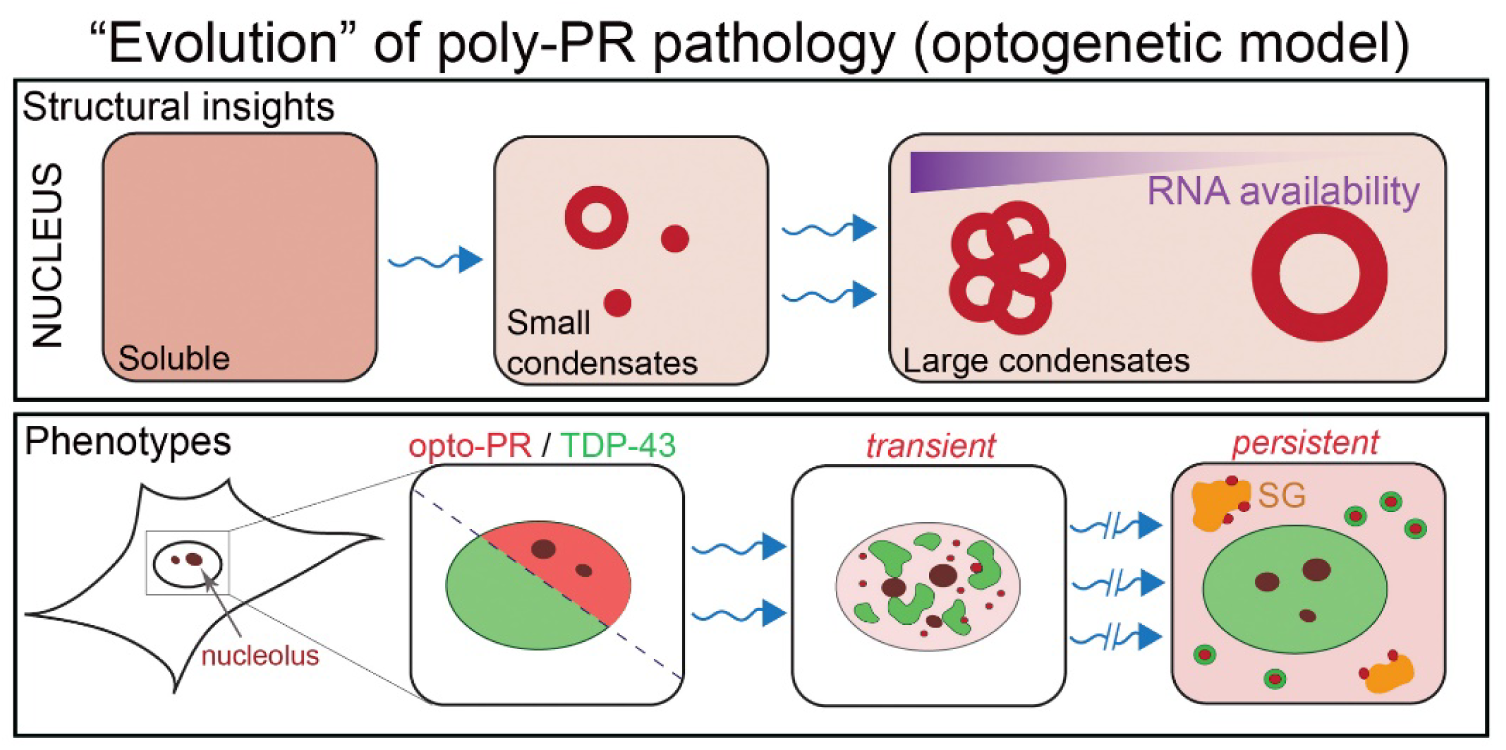

## Introduction

A G_4_C_2_ hexanucleotide repeat expansion (HRE) in the first intron of the *C9orf72* gene is the most common genetic alteration associated with amyotrophic lateral sclerosis (ALS) and frontotemporal dementia (FTD) (DeJesus-Hernandez et al, 2011; Renton et al, 2011). Healthy individuals commonly carry 2 repeats, while C9-ALS/FTD is associated with ≥30 repeat lengths (Van Mossevelde et al, 2017). Production of dipeptide repeat (DPR) proteins from the *C9orf72*-HRE transcripts is one of the proposed mechanisms of repeat toxicity (Ash et al, 2013; Mori et al, 2013; Zu et al, 2013). Both sense and antisense C9orf72-HRE transcripts are translated in all reading frames via non-canonical, repeat-associated non-AUG (RAN) translation to produce five DPRs: poly-GA, -GR (sense), -PR, -PA (antisense) and -GP (both sense and antisense). All five DPRs have been detected in the patient CNS, primarily as cytoplasmic and less often nuclear inclusions in neurons and glia (Gendron *et al*, 2013; Mackenzie *et al*, 2015; Saberi *et al*, 2018; Sakae *et al*, 2018; Schludi *et al*, 2015). Whilst poly-GA and -GP are the most abundant, arginine containing DPRs (R-DPRs) poly-GR and poly-PR are the most toxic species in cells and *in vivo* (Kwon *et al*, 2014; Mizielinska *et al*, 2014; Moens *et al*, 2019). For example, nuclear poly-PR aggregation was sufficient to induce ALS-like phenotypes in non-human primates (Xu *et al*, 2023), and expression of this DPR was extremely toxic in mice (LaClair *et al*, 2020; Zhang *et al*, 2019).

R-DPR inclusion pathology seen in patients and mouse models (Chew *et al*, 2019; Choi *et al*, 2019; Cook *et al*, 2020) is not easily reproducible in cultured cells, where R-DPRs typically display diffuse distribution (outside the nucleolus), even when overexpressed and independent of the repeat length; this is in contrast to poly-GA that readily forms large cytoplasmic aggregates (Frottin *et al*, 2021; Hartmann *et al*, 2018; Liu *et al*, 2022; Lopez-Gonzalez *et al*, 2016; Vanneste *et al*, 2019). High arginine content renders R-DPRs highly hydrophilic and hence soluble. Although poly-PR and poly-GR have similar biochemical properties, molecular dynamics simulations revealed that poly-PR is capable of (limited) self-association – forming either dimers or small amorphous oligomers, whereas poly-GR is not likely to form any stable oligomeric species (Zheng *et al*, 2021). Consistently, *in vitro,* R-DPRs undergo phase separation only in the presence of additional agents – RNA or proteins (Balendra *et al*, 2023; Boeynaems *et al*, 2017; Lee *et al*, 2016; Zhang *et al*., 2019). This suggests that certain factors in the cellular environment drive R-DPR loss of solubility in ALS/FTD. Sensitive immunoassays revealed that soluble DPRs are less abundant in the more affected brain regions, as compared to the relatively spared regions (Quaegebeur *et al*, 2020).

Intermediate products of R-DPR aggregation may contribute to cellular dysfunction, however due to difficulty of modelling this molecular event, their role could not be investigated in the cellular setting. We hypothesised that the use of optogenetics (Park *et al*, 2017; Shin *et al*, 2017) will allow circumventing the intrinsic solubility of R-DPRs, by triggering and maintaining an oligomerised R-DPR state. Indeed, using Cry2olig tagging, we were able to reliably induce condensation of poly-PR (“opto-PR”) in cultured cells, with spatial and temporal control. Using this model, we show that: i) Poly-PR can form nuclear condensates with a specific ordered arrangement reminiscent of TDP-43 “anisosomes” (Yu *et al.,* 2021), and these assemblies trigger abnormal nuclear granulation of TDP-43. ii) Poly-PR cytoplasmic aggregates can form by clustering around stress granules and can nucleate cytoplasmic TDP-43 aggregates, whilst maintaining a TDP-43/DPR demixed state. These findings unveil a converging molecular mechanism for aberrant C9orf72-DPR and TDP-43 species – formation of ordered nuclear condensates. Furthermore, they link putative early disease-stage C9-ALS/FTD species to TDP-43 pathology – the main correlate of neurodegeneration. Finally, they provide a possible mechanism of large DPR inclusion nucleation/seeding in the cytoplasm.

## Results

### Optogenetic modeling of C9orf72 poly-PR condensation in cells

We utilised Cry2olig, a Cry2 variant with high oligomerisation capacity (Taslimi *et al*, 2014), in an attempt to induce and maintain R-DPR self-association. ‘Opto-DPR’ constructs were generated for codon-optimised expression of poly-PR, poly-GR and additionally poly-GP (36 repeats), tagged with Cry2olig and mCherry on the N-terminus (Fig.1A). Opto-DPRs demonstrated a subcellular distribution pattern in HeLa cells typical for the 30-1000 repeat range (Bennion Callister *et al*, 2016; Kanekura *et al*, 2018; Liu *et al*., 2022; Vanneste *et al*., 2019), where poly-GR and -GP were predominantly cytoplasmic and poly-PR was predominantly nuclear, with high enrichment in the nucleolus; none of the DPRs showed signs of aggregation (Fig.1B). Upon single-pulse light stimulation on a custom blue-light array, Cry2olig vector control, opto-PR and opto-GP but not opto-GR readily formed visible clusters/foci (Fig.1B), consistent with the low poly-GR self-association capacity reported in simulation studies (Zheng *et al*., 2021). Opto-PR foci were small (<500 nm), dot-like and exclusively nuclear, whereas opto-GP formed large amorphous cytoplasmic inclusions (Fig.1B). We focused on opto-PR thereafter, using opto-GR as a control.

**Figure 1.**
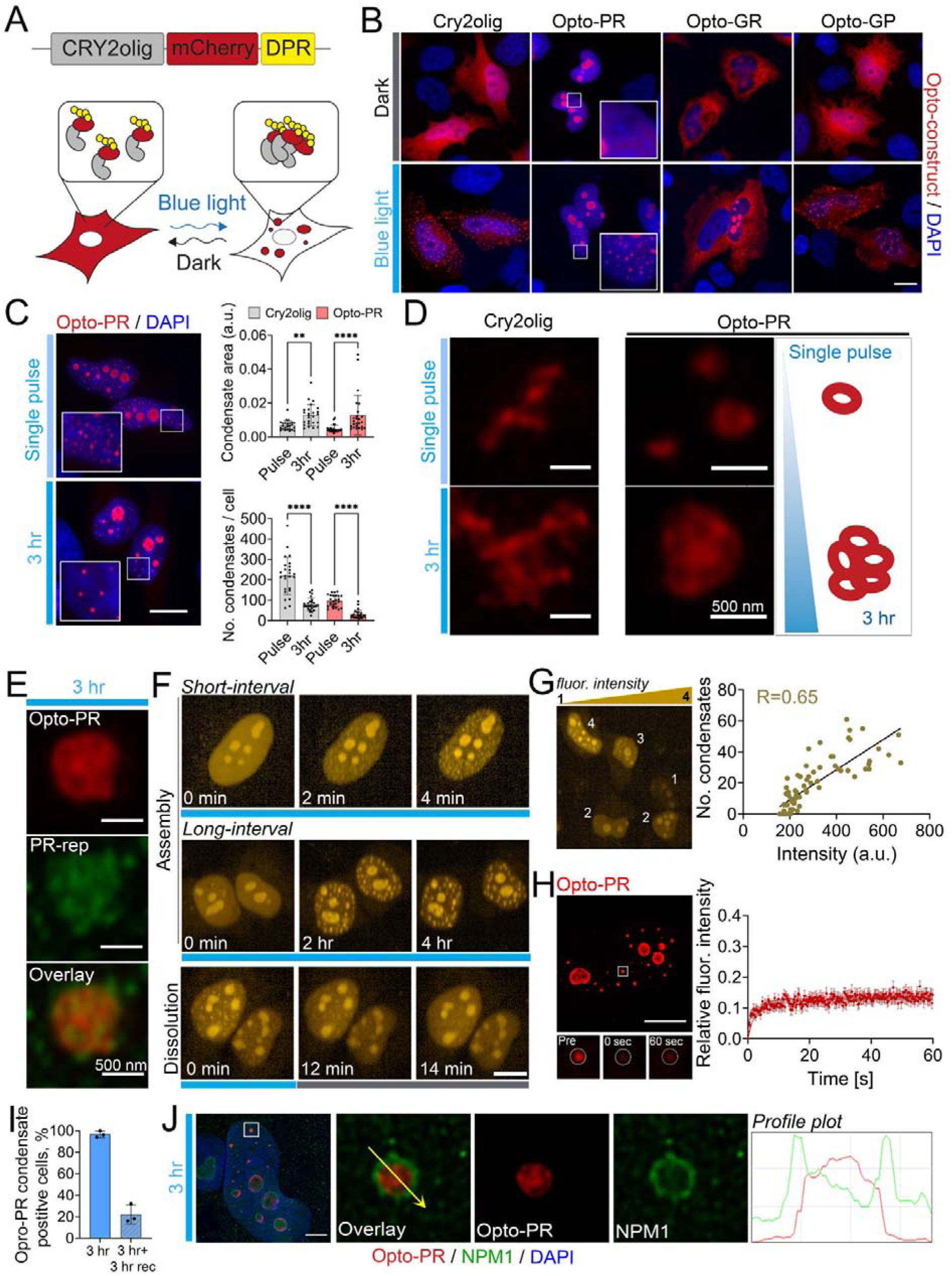
An optogenetic cellular system for controllable *C9orf72* DPR condensation. (A) Opto-DPR approach. (B) Opto-DPR condensation can be induced by a single-pulse blue light stimulation. HeLa cells expressing the respective opto-DPR or empty Cry2olig-mCherry vector were analysed 24 h post-transfection. Blue-light array (single-pulse) was used. Scale bar 10 µm. (C) Continuous opto-stimulation induces larger opto-PR condensates as compared to single light pulse. HeLa cells expressing opto-PR or empty vector were subjected to either a single-pulse or a 3-h continuous blue-light stimulation. 30 cells per condition were analysed from a representative experiment. **p<0.01, ****p<0.0001, Kruskal-Wallis test with Dunn’s post-hoc test. Scale bar 10 µm. (D) Super-resolution microscopy (SRM) demonstrates structural differences between Cry2olig-only and opto-PR assemblies and between the condensates formed after single-pulse and continuous (3-h long) blue-light stimulation. Representative images and graphical representation are shown. (E) Poly-PR, detected using a PR-repeat antibody, is enriched within opto-PR condensate rims. Representative image is shown. (F) Opto-PR condensate induction and tracking using confocal longitudinal imaging. Opto-PR expressing cells were stimulated with a 488 nm laser (at 80% for 500 ms), coupled with mCherry imaging. Cells were stimulated/imaged either every 2 min (“short-interval”) or every 15 min for up to 4 h (“long-interval”). Alternatively, cells with preformed condensates were imaged every 2 min without stimulation (“dissolution”). Representative images are shown. Scale bar, 10 µm. (G) Opto-PR condensate nucleation is concentration-dependent. mCherry fluorescence intensity was measured in the nucleoplasm of individual cells, outside the nucleolus, and the number of condensates was quantified in the same cells at the peak of their assembly (7-min interval repetitive stimulation for 49 min). 75 cells were analysed. Also see Fig.S1C. (H) FRAP analysis after full opto-PR condensate photobleaching reveals low protein mobility between the condensate and nucleoplasm. Representative image and FRAP curve for 25 cells from a representative experiment are shown. Error bars represent SEM. Scale bar, 10 µm. (I) Nuclear opto-PR condensates can become persistent. Opto-PR condensate were induced by 3-h continuous stimulation on blue-light array and the proportion of condensate-positive cells was quantified immediately post-stimulation or after 3 h of recovery in the dark. N=3 (300 cells analysed in total). (J) Opto-PR condensates are positive for nucleophosmin (NPM1). Opto-PR condensates induced by a 3-h continuous opto-stimulation were analysed by SRM. Representative images and profile plots are shown. Scale bar, 2 µm.

Nuclear opto-PR foci/condensates were negative for the two nucleolar markers tested, fibrillarin (FBL) and UPF1, confirming that they are not merely fragments of the nucleolus (Fig.S1A). Continuous 3-h blue-light stimulation led to an increased opto-PR condensate size, as compared to single pulse (Fig.1C). By super-resolution microscopy (SRM), the condensates induced by a single light pulse were found to represent a mixed population of ~100 nm dot-like and ~250 nm spherical, hollow-centre assemblies resembling anisotropic vesicle-like structures formed by arginine-rich peptide/RNA mixtures *in vitro* (Alshareedah *et al*, 2020). Furthermore, the larger condensates formed after 3-h continuous light exposure were found to represent multiples of these spheroids (Fig.1D). Although Cry2olig protein on its own also formed clusters throughout the cell in response to blue light (Fig.1B) (Taslimi *et al*., 2014), these structures appeared disordered/filamentous and were clearly different from opto-PR condensates (Fig.1D). Opto-PR condensates could be detected by a PR-repeat specific antibody used in human tissue studies (Fig.1E). Therefore, poly-PR confers a specific architecture to light-inducible condensates.

We next set up an imaging approach for simultaneous induction and tracking of opto-PR condensation on a confocal high-content imaging system. Even extremely short blue-light exposures and low 488 nm laser power (50 ms/5%) were sufficient to induce opto-PR foci; a combination of 500 ms exposure/80% laser power was used in subsequent experiments as consistently and robustly inducing condensates (Fig.S1B). Visible opto-PR condensates appeared within 2 min post-pulse and could be maintained by both short- and long-interval repetitive opto-stimulation (every 2 min or every 15 min, respectively, Fig.1F). Condensates were reversible, typically resolving within ~14 min after the last light pulse (Fig.1F). Opto-PR condensate nucleation was concentration-dependent, with more structures forming in higher-expressing cells (R=0.65) (Fig.1G; Fig.S1C). FRAP analysis revealed limited dynamics of opto-PR within these assemblies, with low recovery after photobleaching of the entire structure, despite a significant amount of diffuse opto-PR in the nucleoplasm (Fig.1H). A fraction of cells (22.0±8.7%) developed persistent opto-PR condensates after continuous light stimulation, which were still detectable after 3 h of recovery in the dark (Fig.1I).

Poly-PR was previously shown to interact with nucleophosmin (NPM1) and to co-partition with this protein into phase-separated droplets *in vitro* (Lee et al., 2016; White et al, 2019). Consistent with this, opto-PR condensates stained positive for NPM1, where NPM1 formed a “shell” around the condensates, suggesting its secondary recruitment (Fig.1J). Interestingly, opto-stimulation induced opto-PR signal segregation in the nucleolus (Fig.S1D). In this, we observed intra-nucleolar demixing of opto-PR from NPM1, where the proteins fully co-localised in the granular component (GC) under dark conditions but formed two distinct phases after opto-stimulation (Fig.S1E,F). In contrast, opto-PR and FBL (the latter residing in the nucleolar dense fibrillar component, DFC) showed no co-localisation both under dark and light conditions (Fig.S1E,F).

R-DPRs were shown to promiscuously interact with membraneless organelle (MLO) components – RNA and proteins with low-complexity domains, leading to wide-spread MLO dysfunction (Kwon *et al*., 2014; Lee *et al*., 2016; Lin *et al*, 2016; Liu *et al*., 2022). We investigated the effect of opto-PR and its condensation on MLOs, focusing on those in the nucleus due to the predominantly nuclear localisation of this DPR. Systematic analysis of four MLOs – nuclear bodies (Gems, Cajal bodies, paraspeckles and speckles) revealed only minor changes in the number and size in the presence of diffuse or condensed opto-PR (Fig.S2). Cytoplasmic stress granules (SGs) were not induced by opto-PR with or without blue-light stimulation (3 h continuous), consistent with its mainly nuclear localisation (data not shown).

Thus, microscopically visible DPR self-assembly/condensation in cultured cells can be achieved using Cry2olig tagging, allowing the formation of DPR-specific, ordered assemblies. Opto-PR condensates are characterised by concentration-dependent growth and low dynamic properties associated with persistence, and sequester NPM1.

### RNA limits the growth of anisotropic poly-PR condensates in cells

We next asked whether our opto-model can be used to characterise potential modifiers of poly-PR condensation. RNA was previously found to promote R-DPR phase separation *in vitro* (Balendra *et al*., 2023; Boeynaems *et al*., 2017; Gittings *et al*, 2020). More recently, using RNA-protein crosslinking, R-DPRs have been shown to bind RNA in cells, in particular ribosomal RNA (rRNA), with a preference for GA-rich sequences (Balendra *et al*., 2023; Ortega *et al*, 2023). Using electrophoretic mobility shift assay (EMSA) with an RNA oligonucleotide representing a naturally occurring RNA sequence containing 5xGA repeats (Clip34nt) (Bhardwaj *et al*, 2013), we indeed observed that synthetic poly-PR and -GR peptides (10-mers) form complexes with RNA (Fig.2A).

**Figure 2.**
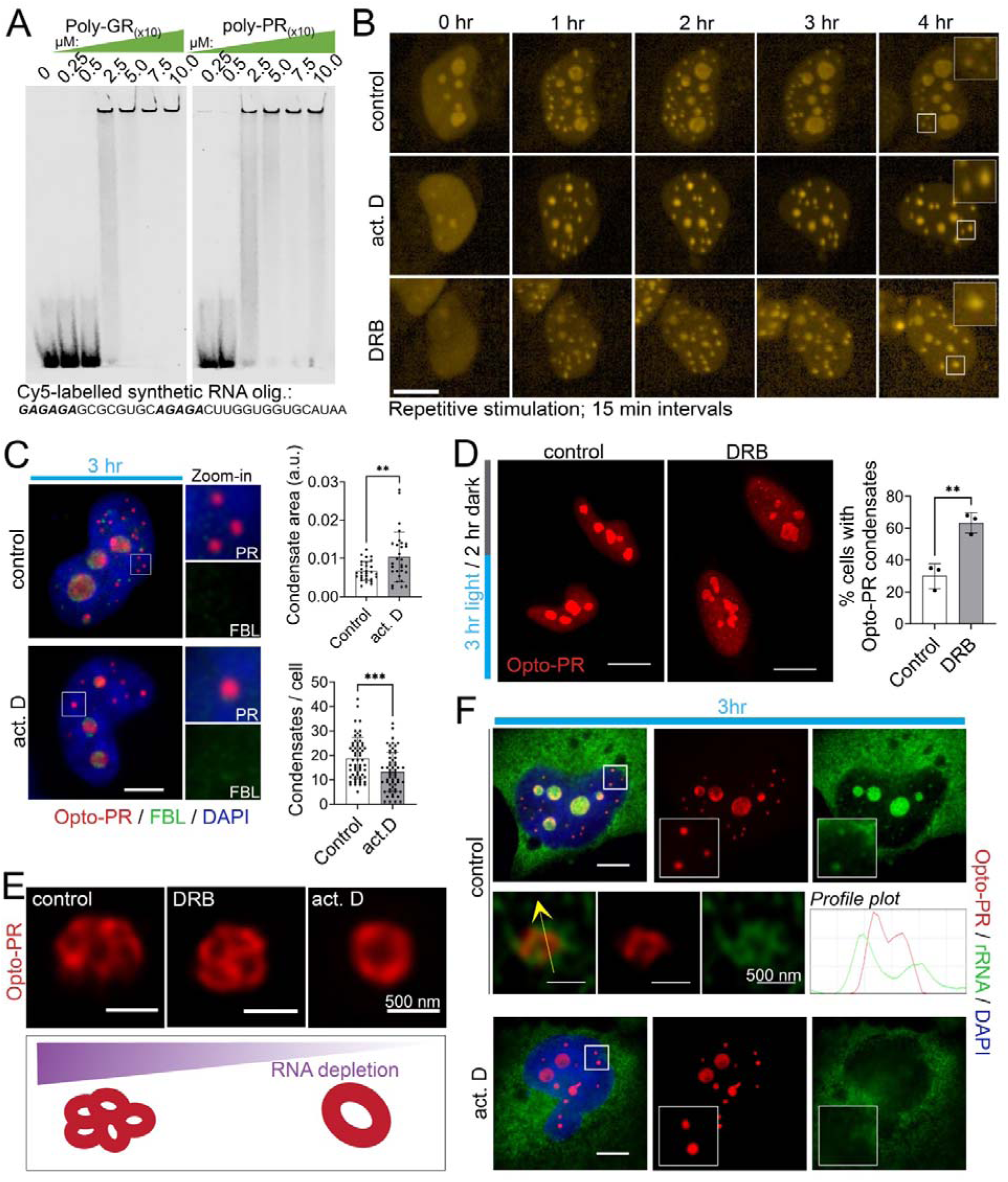
RNA limits opto-PR condensation in the nucleus. (A) Electrophoretic mobility shift assay (EMSA) with a natural GA-rich RNA sequence reveals R-DPR binding to RNA. Cy5-labelled synthetic nucleotide “Clip34nt” (34-mer) and synthetic poly-PR and poly-GR peptides (10 repeats) were used. Representative gel is shown. (B) RNA depletion promotes opto-PR condensate growth. Opto-PR expressing cells were pre-treated with actinomycin D or DRB for 1 h, followed by long-interval repetitive blue-light stimulation (every 15 min) coupled with time-lapse imaging for up to 4 h (in the presence of the inhibitor). Representative images are shown. Scale bar, 10 µm. (C) Opto-PR condensates formed under conditions of actinomycin D-induced RNA depletion are larger in size and less numerous than in RNA-sufficient cells. Cells were opto-stimulated for 3 h continuously. Representative images and quantification are shown. Note that the larger condensates remain FBL-negative. 30 and 60 cells per condition were included in analysis for condensate size and number, respectively, from a representative experiment. **p<0.01, ***p<0.001, Student’s *t* test. Scale bar, 5 µm. (D) Opto-PR condensates formed in RNA-depleted conditions are more persistent. Opto-PR expressing cells were light-stimulated for 3 h continuously, with or without DRB addition, followed by DRB removal and recovery for 2 h in the dark. Representative images and quantification are shown. N=3 (150 cells analysed in total). **p<0.01, Mann-Whitey *U* test. Scale bar, 10 µm. (E) Opto-PR condensates formed in actinomycin D-treated cells are structurally different, as revealed by SRM. Representative images of condensates of a comparable size from control, DRB- or actinomycin D-treated cells induced by 3-h continuous opto-stimulation are shown, alongside with graphical representation. (F) Ribosomal (r)RNA depletion from the nucleus and opto-PR condensates in actinomycin D treated cells. Representative images are shown. Scale bar, 5 µm.

Having confirmed RNA binding, we next studied the impact of RNA depletion on opto-PR condensates. Opto-PR expressing cells were treated with two transcriptional blockers, a global inhibitor (actinomycin D) and an RNA polymerase II-specific inhibitor (5,6-Dichloro-1-β-D-ribofuranosylbenzimidazole, DRB), followed by induction and tracking of opto-PR condensates in individual cells for up to 4 h (long-interval repetitive stimulation, in the presence of the inhibitor). Both inhibitors caused dramatic nucleolar shrinking confirming their activity in cells (Fig.2B) (Shav-Tal *et al*, 2005). Opto-PR condensates formed in RNA-depleted conditions appeared noticeably larger compared to control (Fig.2B). Similar result was obtained with a 3-h continuous opto-stimulation on blue-light array, and quantification confirmed a significant increase in the opto-PR condensate size (Fig.2C). These larger structures remained negative for the nucleolar marker FBL (Fig.2C). DRB is a reversible inhibitor, which allowed us to analyse possible changes in the stability of RNA-depleted opto-PR condensates. Cells were stimulated for 3 h in the presence or absence of DRB followed by 2 h of recovery in the dark. Opto-PR condensates formed in the presence of DRB appeared significantly more persisting, still detectable following the recovery in 63±6.4% cells, compared to 30±7.8% cells in control condition (Fig.2D).

Increase in the opto-PR condensate size upon actinomycin D treatment was accompanied by a decrease in their number (Fig.2C), suggesting clustering or fusion. SRM analysis revealed that opto-PR condensates in actinomycin D treated cells were no longer clusters of individual ~250 nm spherical units seen in the untreated or DRB condition but instead represented larger (>500 nm) hollow-centre spheres (Fig.2E). RNA on the surface of condensates has been shown to limit their fusion, including in cells (Cochard *et al*, 2022). Actinomycin D but not DRB depletes ribosomal RNA (rRNA), and we observed dense rRNA signal on the opto-PR condensate surface which disappeared after actinomycin D treatment (Fig.2F). Therefore, the opto-PR condensates in actinomycin D treated cells may form by fusion of the smaller units due rRNA depletion from the surface, followed by relaxation into a larger spherical assembly.

Another putative modifier of R-DPR self-association/condensation is arginine dimethylation (DMA), which increases the fluidity of R-DPR droplets *in vitro* and was found enriched in R-DPR inclusions in C9-ALS/FTD patient tissue (Gittings *et al*., 2020). We employed two small molecule methyltransferase inhibitors, MS023 inhibiting five type-I methyltransferases that synthesise asymmetric DMA (aDMA) and EPZ015666, a specific inhibitor of PRMT5 responsible for most symmetric DMA (sDMA) (Chan-Penebre *et al*, 2015; Eram *et al*, 2016). Opto-PR expressing cells were treated with MS023 and EPZ015666 for 24 h and then exposed to blue-light for 3 h continuously, followed by opto-PR condensate quantification. We observed a mild decrease in the condensate number after aDMA depletion without changes in their size or structure (Fig.S3). Thus, removing DMA marks may attenuate opto-PR condensate nucleation, although this effect is small.

Therefore, our cellular opto-PR model can be utilised to analyse modifiers of poly-PR self-assembly in the cellular context, as exemplified by the modulatory effect of RNA that we have uncovered.

### Poly-PR condensation induces nuclear TDP-43 pathology

R-DPR interactomes are enriched in RNA-binding proteins (RBPs) (Kwon *et al*., 2014; Lee *et al*., 2016). We therefore examined the effect of nuclear opto-PR condensation on ALS/FTD relevant RBPs. TDP-43, FUS, NONO and SFPQ, all tagged with GFP, were co-expressed with opto-PR and their subcellular distribution was examined with and without opto-stimulation. Opto-PR presence *per se* did not affect RBP distribution (Fig.3A; Fig.S4A). However, light-stimulated opto-PR expressing cells displayed a striking nuclear condensation phenotype for TDP-43 but not other RBPs analysed (Fig.3A; Fig.S4A). Although we did observe TDP-43 condensation in a fraction of light-stimulated Cry2olig-expressing, it was significantly smaller than in opto-PR cultures; nor was it observed in light-stimulated opto-GR expressing cells (Fig.3B). Opto-PR assemblies were frequently found in the physical contact with TDP-43 condensates (44±0.5% of all opto-PR foci) (Fig.3C), suggestive of a direct nucleating effect of oligomerising opto-PR. Fractionation confirmed reduced solubility of TDP-43 GFP after induction of opto-PR condensation but not in light-stimulated opto-GR expressing cells (Fig.3D). This analysis also confirmed reduced solubility of opto-PR but not opto-GR in light-stimulated cells (Fig.3D).

**Figure 3.**
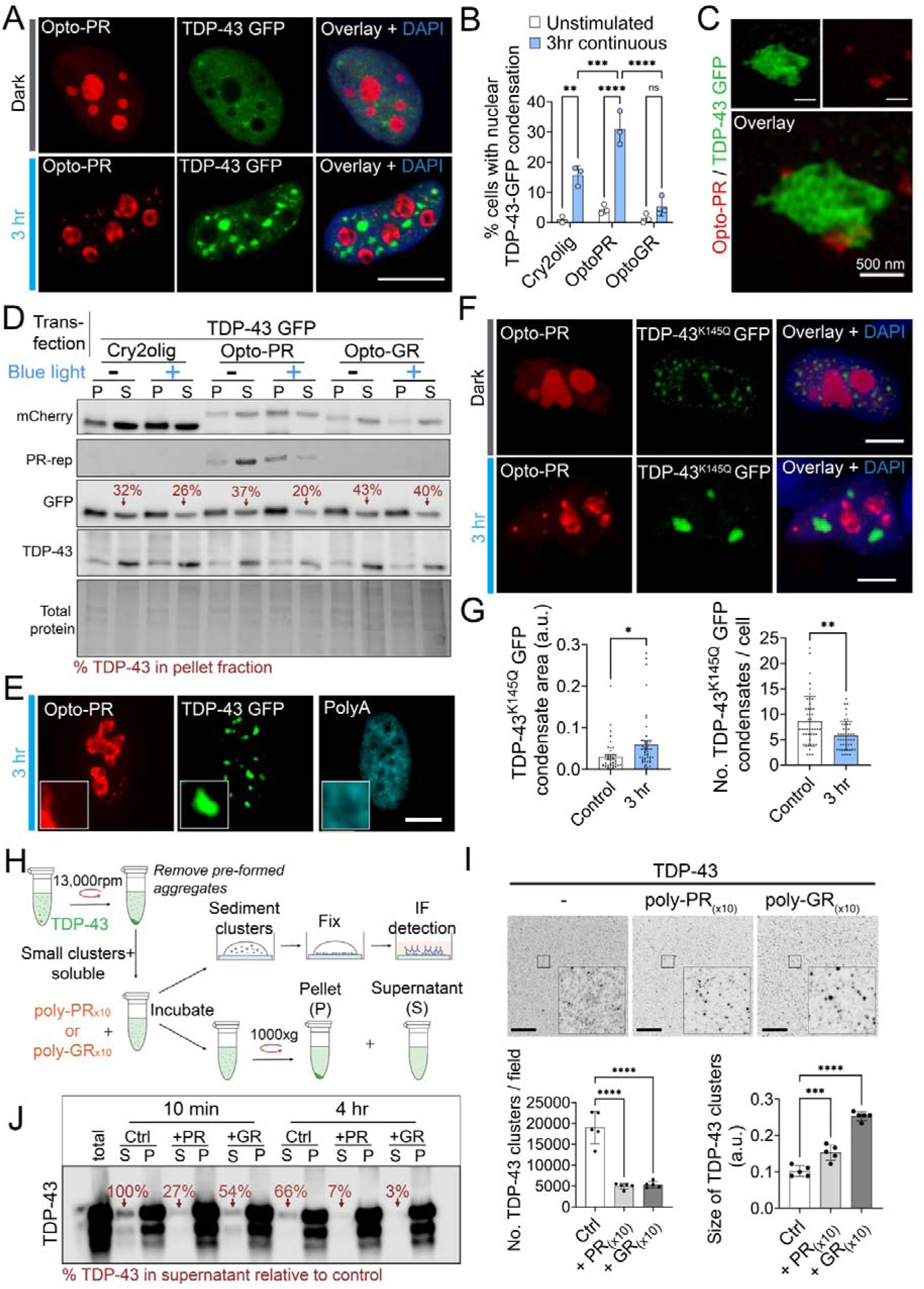
Nuclear opto-PR condensation induces TDP-43 pathology. (A,B) Opto-PR self-assembly induces nuclear condensation of co-expressed TDP-43 GFP. Cells were opto-stimulated for 3 h continuously. Representative images (A) and quantification (B) are shown. Scale bar, 10 µm. N=3 (≥100 cells analysed per biological repeat). **p<0.01, ***p<0.001, mixed-effects analysis with Sidak’s multiple comparisons test. (C) Spatial proximity of opto-PR and TDP-43 GFP nuclear condensates as revealed by SRM. Representative image is shown. (D) Reduced TDP-43 GFP solubility upon induction of opto-PR condensation. Cells expressing opto-PR were subjected to 3-h continuous blue-light stimulation before lysis and fractionation. Western blots and quantification of the relative TDP-43 GFP amount in the supernatant fraction are shown for a representative experiment. S, supernatant; P, pellet. (E) TDP-43 condensates induced by opto-PR self-assembly are devoid of polyA+ RNA. Cells after 3-h continuous opto-stimulation were analysed by poly(dT) RNA-FISH. Representative image is shown. Scale bar, 5 µm. (F,G) Opto-PR self-assembly promotes nuclear condensation of a TDP-43 acetylation mimic mutant (K145Q). Representative images (F) and quantification (G) are shown. 50 cells were analysed from a representative experiment. *p<0.05, **p<0.01, Student’s *t* test. Scale bar, 5 µm. (H-J) R-DPRs promote TDP-43 clustering *in vitro.* Recombinant TDP-43 (supernatant fraction depleted of large aggregates; 1 µM) was incubated with equimolar amounts of poly-PR_(x10)_ or poly-GR_(x10)_ peptides and analysed by immunostaining and fractionation/western blot. Methodology (H), immunostaining/imaging (I) and fractionation/western blotting (J) data are shown. In I, 10 min incubation was used and five fields of view from a representative experiment were analysed; ***p<0.001, ****p<0.0001, one-way ANOVA with Dunnett’s post-hoc test. In J, representative western blot and band intensity quantification for the supernatant are shown. S, supernatant; P, pellet. Scale bar, 20 µm.

TDP-43 GFP nuclear condensates induced by opto-PR were strikingly similar in their morphology to the condensates formed during the recovery from arsenite stress (Wang *et al*., 2020; Cohen *et al*., 2015; Huang, Ellis *et al*., manuscript in revision). Like these stress-induced foci, TDP-43 condensates induced by opto-PR were devoid of polyA+ RNA (Fig.3E). We therefore asked whether TDP-43 condensation elicited by opto-PR was due to an upregulated stress response. Classic cellular stress markers, phosho-eIF2α, GADD34 and ATF4, were not altered in opto-PR expressing cells after 3-h light stimulation (Fig.S4B,C). This is in contrast to the dramatic upregulation of these markers during the recovery from sodium arsenite stress used as a positive control (Fig.S4C). In addition to demonstrating the stress-unrelated nature of TDP-43 condensates induced by opto-PR self-assembly, this experiment also confirmed a lack of phototoxicity in our model.

TDP-43 acetylation was shown to impair its RNA binding and enhance aggregation, with acetylated TDP-43 inclusions detected in sALS (Cohen *et al*, 2015). We investigated the effect of opto-PR condensation on an acetylation-mimic TDP-43 mutant, K145Q (Cohen *et al*., 2015). In agreement with the published data, TDP-43 K145Q was prone to spontaneously forming nuclear foci/granules in naïve cells (Fig.S4D). Opto-PR condensation potentiated the pro-aggregating effect of acetylation, further increasing nuclear granulation of TDP-43 K145Q (Fig.3F,G).

We next examined the effect of R-DPRs on TDP-43 higher-order assembly *in vitro,* using the condensate immunodetection/imaging assay we recently developed (Hodgson, Huang *et al.,* 2024, doi: 10.2139/ssrn.4721338) (Fig.3H). Both poly-PR and poly-GR peptides were included in these studies. “Supernatant” fraction of recombinant TDP-43 (after removal of preformed aggregates) containing small protein clusters and soluble protein was incubated with equimolar amounts of synthetic R-DPR peptides, followed by sedimentation and fixation of TDP-43 clusters on coverslips for immunostaining/imaging; in parallel, samples were fractionated by centrifugation for western blot analysis (Fig.3H). Addition of both R-DPRs significantly enhanced TDP-43 clustering – manifested as an increased cluster size and decreased cluster number, with poly-GR being more potent (Fig. 3I). Fractionation confirmed increased partitioning of TDP-43 to the pellet fraction in the presence of R-DPRs (Fig.3J). In contrast, addition of a peptide with a “generic” sequence moderately enriched in proline and containing no arginine residues (V5: GKPIPNPLLGLDST) did not induce TDP-43 clustering (Fig.S4E), ruling out a non-specific molecular crowding effect of R-DPRs.

Collectively, these results suggest that poly-PR condensation can directly cause changes to the nuclear distribution of TDP-43 without activation of stress signaling.

### Cytoplasmic poly-PR assemblies are persistent and selectively sequester TDP-43

Having characterised the nuclear phenotypes, we asked whether our opto-PR model is amenable to reproducing the cytoplasmic pathology typical for DPRs. Continuous 24-h long stimulation on the blue-light array resulted in significant redistribution of opto-PR to the cytoplasm, with cytoplasmic foci formation in 32% of cells (>300 transfected cells analysed; Fig.4A,B). This was accompanied by a reduction in the incidence of nuclear opto-PR condensates (from 94% cells after 3 h to 9% after 24 h of stimulation) (Fig.4A,B). Nuclear and cytoplasmic opto-PR foci induced by 24-h opto-stimulation were persisting, with no significant decline observed after 8 h of recovery in the dark (Fig.4A,B). This is in contrast to the nuclear opto-PR condensates forming after a 3-h stimulation that were largely cleared after 3 h of recovery in the dark (Fig.4A; Fig.1I). Furthermore, we found that in a small proportion of cells that developed spontaneous SGs, opto-PR assemblies surrounded SGs (Fig.4C). Opto-PR redistribution to the cytoplasm was not due to the nuclear membrane damage/nuclear pore complex disruption, since nuclear retention of several RBPs was not affected in these cells (Fig.4D). Therefore, impaired nuclear import of opto-PR under these conditions could be due to its submicroscopic oligomerisation in the cytoplasm. Strikingly, endogenous TDP-43 but not other ALS-related RBPs (FUS, NONO, SFPQ) were found to be enriched in the cytoplasmic opto-PR foci (Fig.4D). In contrast, Cry2olig-only cytoplasmic structures induced by 24-h light stimulation were negative for TDP-43 (Fig.4E). SRM revealed that cytoplasmic opto-PR assemblies were ~250 nm structures that, unlike nuclear condensates, were not hollow (Fig.4F). It also revealed that in these assemblies, opto-PR and TDP-43 remained demixed, with the TDP-43 signal primarily on the surface (Fig.4F). A small fraction of cytoplasmic opto-PR foci were positive for p62 (Fig.4G) but none of them stained positive for ubiquitin (data not shown).

**Figure 4.**
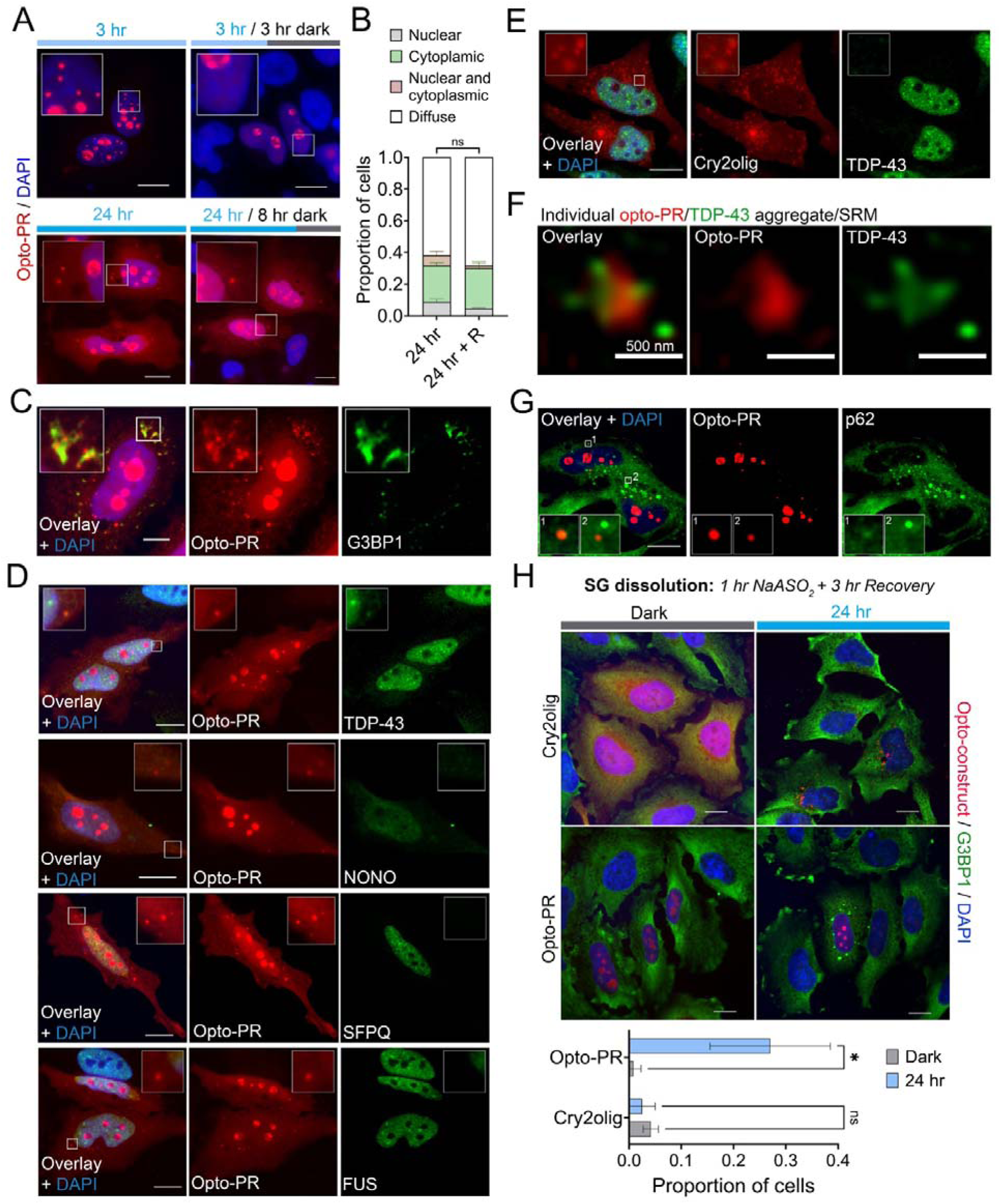
Prolonged opto-stimulation elicits cytoplasmic opto-PR/TDP-43 pathology. (A,B) Prolonged, 24-h blue-light stimulation leads to cytoplasmic opto-PR redistribution and aggregation, with persistent assembly formation. Proportion of cells with opto-PR assemblies (nuclear and cytoplasmic) was quantified after 24-h blue-light array stimulation with and without 8-h recovery in the dark. Representative images (A) and quantification (B) are shown. Images for a 3-h stimulation and recovery are included for comparison. N=3 (300 cells analysed per condition). n.s., non-significant. Scale bar, 10 µm. (C) Cytoplasmic opto-DPR foci surround spontaneous stress granules (SGs, visualised using G3BP1 as a marker). Representative image is shown. Cells were opto-stimulated for 24 h continuously. Scale bar, 5 µm. (D) Endogenous TDP-43 but, not other RBPs, joins opto-PR assemblies induced by prolonged opto-stimulation. Also note normal nuclear localisation of all RBPs in cells with cytoplasmic opto-PR. Cells were opto-stimulated for 24 h continuously. Representative images are shown. Scale bars, 10 µm. (E) Cry2only cytoplasmic assemblies are negative for TDP-43. Cells were opto-stimulated for 24 h continuously. Representative images are shown. Scale bars, 10 µm. (F) TDP-43 remains demixed within opto-PR within cytoplasmic assemblies, as revealed by SRM. Cells were opto-stimulated for 24 h continuously. Representative image is shown. (G) Cytoplasmic opto-PR assemblies are occasionally positive for p62. Representative image is shown. Insets 1 and 2 shows examples of p62-positive and -negative assemblies, respectively. Scale bars, 10 µm. (H) SG dissolution is affected in cells with cytoplasmically localised opto-PR. SGs Representative were induced NaAsO_2_ in cells expressing either opto-PR or Cry2olig-only, subjected to 24-h long opto-stimulation or kept in the dark. Efficiency of SG dissolution was analysed 3 h into the recovery (post-NaAsO_2_ removal). Representative images and quantification are shown. N=3 (120 cells analysed per condition). *p<0.05, Two-way ANOVA with Sidak’s multiple comparisons test. Scale bars, 10 µm.

We next examined whether opto-PR accumulated/aggregated in the cytoplasm after prolonged opto-stimulation affects stress-induced SGs (Boeynaems *et al*., 2017; Marmor-Kollet *et al*, 2020; Wen *et al*, 2014). Opto-PR and Cry2olig-only expressing cells subjected to long opto-stimulation or kept in the dark were analysed during the recovery from NaAsO_2_ (3 h time-point), using G3BP1 as a marker. SG disassembly was delayed in light-stimulated opto-PR expressing cells compared to unstimulated opto-PR cells or Cry2-only expressing (stimulated or unstimulated) cells (Fig.4H). This finding further validates our cellular model as capable of reproducing the key molecular effects of R-DPRs on the cellular RNA/RNP granule metabolism.

Therefore, our opto-model is amenable to the induction of cytoplasmic poly-PR accumulation and aggregation, with its cytoplasmic assemblies being significantly more persistent than those in the nucleus. Our data also point to a role for SGs in the growth of cytoplasmic poly-PR aggregates as well as a role for poly-PR assemblies in “nucleating” TDP-43 aggregation in the cytoplasm.

## Discussion

A vast body of knowledge on C9-DPR related disease mechanisms has accumulated in the past decade yet it remains unclear whether, and if so how, DPR self-assembly, resulting in a C9-ALS/FTD hallmark pathology, contributes to the disease. Aggregation intermediates, and especially smaller, highly reactive oligomeric species, have been validated as toxic/pathogenic in the case of other neurodegeneration-linked proteins, such as tau and alpha-synuclein (Bengoa-Vergniory *et al*, 2017; Choi & Gandhi, 2018; Jucker & Walker, 2018). It is possible that equivalent DPR aggregation products play a role in C9-ALS/FTD. Poly-PR is the least abundant DPR in human tissue (Mackenzie et al., 2015; Davidson et al., 2016) despite theoretically it should be expressed at the levels similar to other DPRs (except poly-GP produced from both strands). Although this can be due to a variation in the expression mechanisms and antibody detection, it is also possible that neurons that accumulate poly-PR are lost early in disease due to its high toxicity, including its aggregation products. Although the inclusions of all five DPRs in the patient CNS are morphologically similar (Gendron *et al*., 2013), DPRs other than GA fail to form microscopically visible assemblies in cell models (Frottin *et al*., 2021; Liu *et al*., 2022; Zhou *et al*, 2017). In order to circumvent the high solubility of R-DPRs in cells, we harnessed the Cry2olig opto-module (Taslimi *et al*., 2014) successfully used previously to promote self-association of neurodegeneration-linked proteins (Berard *et al*, 2022; Jiang *et al*, 2021; Mann *et al*, 2019). In line with the molecular dynamics predictions (Zheng *et al*., 2021), poly-GR’s self-association capacity was too low even when facilitated by Cry2olig, not yielding visible condensation in cells. However we succeeded in achieving the condensation of the oligomerisation-competent poly-PR.

Two key phenotypes – alterations to the nucleolus and SGs – validate our opto-PR model in the context of the existing literature (Kwon *et al*., 2014; Marmor-Kollet *et al*., 2020; White *et al*., 2019). In our model, SG disassembly was impaired by cytoplasmic opto-PR (but not Cry2olig-only). This can be caused by altered composition, and hence dynamics, of SGs formed in the opto-PR rich milieu. We also observed sequestration of NPM1, a confirmed R-DPR interactor, into nuclear opto-PR condensates. Beyond these two phenotypes however, we failed to detect the ubiquitous MLO disruption by poly-PR reported in some studies, and its condensation also had limited effect on MLOs. This may be due to high variability of these phenotypes depending on the cell type, DPR levels and repeat length. It would be important to establish consensus phenotypes that can be used for the benchmarking of novel cellular DPR models, and the nucleolar and SG pathology validated in multiple studies are the prime candidate readouts.

We found that poly-PR confers a specific ordered arrangement to the opto-induced nuclear condensates – sphere with hollow centre. Arginine-rich peptides form such anisotropic structures in the presence of RNA *in vitro*, both under RNA and peptide excess conditions (Alshareedah *et al*., 2020). Adopting this model, we speculate that upon interaction with cellular RNA, the small opto-PR oligomers nucleated with the aid of Cry2olig, form nanocondensates with a neutral “head” and a charged “tail” that subsequently coalesce into RNA-coated micelles (~100 nm granules). These transition into a vesicle-like conformation “layered” with RNA on both the internal and external surface (~250 nm granules). Upon (r)RNA depletion, these condensates undergo fusion into a larger (>500 nm) hollow-centre structure (Fig.S5A). Anisotropic condensate is a non-equilibrium state that may require ATP to be established and maintained (Bergmann *et al*, 2023), and in our system, this state is fuelled by the light-induced Cry2olig oligomerisation. Poly-PR was shown to form nuclear structures in C9-ALS/FTD, varying from compact inclusions to less dense “territories”, in multiple studies (Cooper-Knock et al., 2015; Mori et al., 2013; Davidson et al., 2016; Wen et al., 2014). Nuclear poly-PR condensates not overlapping with nucleolar markers also form in transgenic mice (Zhang *et al*., 2019). It would be interesting to establish whether the structures seen in mice adopt a hollow-centre structure during their biogenesis. Anisotropic nuclear assemblies (“anisosomes”) are formed by acetylated TDP-43 in cultured cells and *in vivo* (Yu *et al*, 2021). RBPs complexed with polyadenylated RNA form similar structures in cell models of spinal muscular atrophy (Narcis *et al*, 2018). Furthermore, nuclear condensates of RNA-binding deficient TDP-43 and ALS-linked CREST protein (Kukharsky *et al*, 2015) also possess this typical morphology (Fig.S5B). Remarkably, DDX3X mutations causative of neurodevelopmental disorders also form cytoplasmic hollow condensates, and those composed of an aggressive RNA-binding deficient mutant display low recovery in FRAP (Owens *et al*, 2023). It is possible that specific changes to the cellular metabolism in neurological disease, e.g. altered protein and RNA stoichiometries, favour this assembly type. RNA facilitates LLPS of poly-PR *in vitro* (Balendra et al., 2023) however cellular RNA restricts the growth of the non-dynamic opto-PR condensates in cells. Together with the reports on the solubilising effect of RNA on RBPs (Grese *et al*, 2021; Maharana *et al*, 2018; Mann *et al*., 2019; Shelkovnikova *et al*, 2014a) and on the wide-spread RNA degradation in ALS (Tank *et al*, 2018), our findings suggest than declining RNA levels may be a common factor underlying protein aggregation across ALS subtypes including C9-ALS.

Transient nuclear TDP-43 condensation is a hallmark of stress response (Cohen *et al*., 2015; Wang *et al*., 2020). We recently found that this molecular event leads to TDP-43 loss of function and prolonged STMN2 depletion and is dysregulated by TDP-43 mutations (Huang, Ellis *et al.,* 2024, manuscript in revision). These phenotypes may become persistent with chronic or repetitive stress, precipitating the disease. Here we show that poly-PR self-assembly causes TDP-43 condensation in the absence of stress signalling, indicating the convergence between C9-ALS and other ALS subtypes in this putative disease mechanism. Consistently, recent use of a specific RNA aptamer revealed abnormal nuclear TDP-43 granulation in motor neurons in ALS tissue (Spence et al., 2024).

Cytoplasmic poly-PR assemblies in our opto-model are structurally dissimilar to and more persistent than the nuclear condensates, probably reflecting a different environment in the two cellular compartments. This higher stability is consistent with a higher frequency of poly-PR cytoplasmic inclusions in patients (Zu *et al*., 2013). DPR inclusions are rarely found to co-deposit with TDP-43 inclusions in the patient tissue, with clear anatomical region specificity (Al-Sarraj et al, 2011; Davidson et al., 2016). Some histopathological studies found evidence of DPR aggregation preceding the cytoplasmic TDP-43 pathology (Baborie *et al*, 2015; Vatsavayai *et al*, 2016). TDP-43 joins the cytoplasmic opto-PR foci where the two proteins remained in two phases. It is tempting to speculate that transient cytoplasmic poly-PR assemblies “seed” TDP-43 pathology. SGs may play an equivalent role for cytoplasmic poly-PR inclusions, concentrating the initial poly-PR assemblies in a confined space and promoting their coalescence into a larger aggregate. Indeed, a recent study showed that cytoplasmic TDP-43 aggregates can be nucleated within SGs, as a separate phase, and left behind after SG dissolution (Yan *et al*, 2024). Inclusions C9-ALS/FTD are likely composed of several DPRs, and their co-expression leads to different phenotypes *in vivo* (West *et al*., 2020). Opto-GP readily undergoes cytoplasmic condensation in our model, enabling future opto-DPR co-aggregation studies.

### Limitations of the study

We used a relatively short repeat length (which nevertheless is in the patient range), whereas the Cry2olig-mCherry is a large tag. However, multiple studies have demonstrated that DPR repeat lengths of 30-1000 have identical subcellular localisation with similar toxicity (Bennion Callister *et al*., 2016; Kwon *et al*., 2014; Miyagi *et al*, 2023; White *et al*., 2019). We cannot exclude that longer repeats will undergo condensation more readily, which should be tested in future studies. In addition, for this proof-of-principle study, non-neuronal cells were used for the ease of expression and imaging. To address potential cell-specificity of the phenotypes revealed in this study, the opto-PR model should be transferred into a (human) neuronal system.

## Method details

### Plasmids

The CRY2olig-mCherry backbone vector was a gift from Chandra Tucker (Addgene plasmid # 60032) (Taslimi *et al*., 2014). DPR plasmids were a gift from Kurt De Vos (Bauer *et al*, 2022). Codon-optimised DPR sequences were cloned into the CRY2olig-mCherry vector using standard techniques. These plasmids are available via Addgene. Plasmids for the expression of GFP-tagged TDP-43, FUS, NONO, SFPQ and CREST (N-terminal tag, in pEGFP-C1 vector) were generated previously (Kukharsky *et al*., 2015; Shelkovnikova *et al*, 2014b). TDP-43 K145Q and F147/149L in pEGFP-C1 vector were generated using PCR mutagenesis.

### Cell culture, transfection and treatments

HeLa cells were obtained from ATCC via Sigma, cultured in Dulbecco’s Modified Eagle Medium/Nutrient Mixture F-12 (DMEM/F-12) supplemented with 10% foetal bovine serum (FBS) and penicillin-streptomycin. For time-lapse imaging, cells were plated on PhenoPlate-96 (black, optically clear bottom, PerkinElmer) at a density of 2 x 10^4^. For all other experiments, cells were seeded in 24 well plates, with or without coverslips dependent on the application, at a density of 5 x 10^4^ cells, unless otherwise stated. Transfection was performed 24 h prior to blue light stimulation, using either Lipofectamine 2000 (ThermoFisher) or jetPRIME (Jena Bioscience) according to the manufacturer’s instructions. For transcriptional inhibition cells were treated with 2.5 µg/ml actinomycin D or 5,6-dichloro-1-beta-D-ribofuranosylbenzimidazole (DRB) (both Sigma). For inhibition of arginine methylation, cells were treated with 10 µM MS-023 or EPZ015666 (both ApexBio). To induce cellular stress, cells were treated with 250 µM NaAsO_2_ (Sigma). Incubation times for treatments within individual experiments are indicated in respective Figure legends.

### Opto-stimulation

Cells expressing opto-constructs were stimulated with a 488 nm laser on Opera Phenix HCS (500 ms, 80% laser power) for live cell time-lapse imaging (under full environmental control conditions), or on a custom-built LED blue-light array housed in a humidified incubator maintained at 37°C with 5 % CO_2_. Cells were protected from light between experiments and during fixation.

### Immunocytochemistry and RNA-FISH

Immunocytochemistry was performed as described elsewhere (Shelkovnikova *et al*, 2018) using the following commercially available antibodies: FBL (rabbit polyclonal, Proteintech, 16021-1-AP); NPM1 (mouse monoclonal, Proteintech, 60096-1-Ig); UPF1 (rabbit polyclonal, Proteintech, 23379-1-AP); PNN (rabbit polyclonal, Proteintech, 18266-1-AP); coilin p80 (mouse monoclonal, BD Biosciences, 612074); SMN (mouse monoclonal, BD Biosciences, 610647); G3BP1 (rabbit polyclonal, Proteintech, 13057-2-AP); repeat-PR (rabbit polyclonal, Proteintech, 23979-1-AP); TDP-43 (rabbit polyclonal C-terminal, Sigma); FUS (mouse monoclonal, Santa Cruz, [4H11], sc-47711); NONO (rabbit polyclonal, Proteintech, 11058-1-AP); SFPQ (rabbit monoclonal, Abcam, ab177149); ribosomal RNA (mouse monoclonal Y10b, NB100-662); p62/SQSTM1 (mouse monoclonal, MAB8028R, R&D Biosystems); RNA-FISH using a Cy5-labelled dT_(30)_ DNA oligonucleotide probe (Sigma) was performed as described earlier (Shelkovnikova *et al*, 2017).

### Microscopy

Conventional fluorescence microscopy was performed using 100x objective on an Olympus BX57 upright microscope equipped with an ORCA-Flash 4.0 camera (Hamamatsu) and cellSens Dimension software (Olympus). Super-resolution microscopy was performed using a 63x oil immersion objective on a ZEISS 980 laser scanning confocal microscope (LSM) with Airyscan 2 detector and ZEN Blue software. Time-lapse microscopy was performed using a 40x objective on Opera Phenix HCS, and Harmony 4.9 software was used for image processing and analysis (all PerkinElmer). Image processing and profile drawing was done using Image J or ZEN Blue software. Condensate/aggregate quantification was done manually or on Image J in a blinded manner.

### Fluorescent recovery after photobleaching (FRAP)

Cells seeded at a density of 2.8 x 10^5^ in glass-bottomed 35mm dishes (Ibidi), were transfected and 24 h post-transfection, subjected to 3-hr continuous stimulation prior to FRAP analysis. Imaging was performed using a 63x oil immersion objective on a ZEISS LSM 800 confocal microscope, equipped with a humidified incubation chamber maintained at 37°C with 5 % CO_2_. FRAP acquisition was performed on condensates formed after 3 h of continuous stimulation. A circular region of interest (ROI) around each condensate was bleached using a 568 nm laser at 100 % laser power. Images were captured pre-bleach, immediately following bleach and at x 200 ms intervals during recovery. The mean fluorescence intensity within the ROI was determined for each image using ZEN blue software. Intensity values were corrected for bleaching during imaging and normalised to pre-bleach intensity. Average values were plotted and FRAP curves fitted using a one-phase association equation in GraphPad Prism 9 software.

### Electrophoretic mobility shift assay (EMSA)

Cy5-labeled RNA oligonucleotide 5’-GAGAGAGCGCGUGCAGAGACUUGGUGGUGCAUAA-3’ (“Clip34nt”) (Bhardwaj *et al*., 2013) was custom-synthesised by Eurofins. Poly-GR_x10_ and poly-PR_x10_ were custom-made by Biosynth. Lyophilised peptides were resuspended in water at 50 mM and stored at −80°C before use. RNA was incubated at 250nM with 250nM-10µM peptides in the EMSA buffer (50 mM Tris-HCl, pH 7.5, 100 mM KCl, 2 mM MgCl_2_, 100 mM β-mercaptoethanol, 0.1 mg/ml BSA) for 15 min at RT with gentle shaking. Samples were analysed on 6% native acrylamide gel in TBE buffer, followed by imaging on Licor Odissey FC (700nm channel).

### Cell lysis, fractionation, and western blotting

For fractionation, cells were lysed in 1 % Titon-X-100 in PBS for 5 min at RT. Lysates were incubated on ice for 30 minutes with vortexing at 5 min intervals and centrifuged at 13,000rpm for 20 min at 4°C. Supernatant and pellet were mixed with 2x Laemmli buffer (2xLB), vortexed and heated at 95°C for 10 min. For western blotting, proteins were resolved on a 10% Mini-PROTEAN® TGX™ hand-cast protein gel and transferred to a PVDF membrane. Gels were stained with Gelcode^TM^ (ThermoFisher) post-transfer for total protein. Membranes were blocked for 1 h in 4 % milk/TBST and then incubated with the following primary antibodies (1:1000 dilution) at 4°C overnight: mCherry (rabbit polyclonal, Proteintech, 26765-1-AP); GFP (mouse monoclonal, 66002-1-Ig, Proteintech); TDP-43 (rabbit polyclonal, C-terminal, Sigma); eIF2α (rabbit polyclonal, Cell Signaling, 9722), eIF2αP (rabbit polyclonal, Abcam, antibody [E90], ab32157); repeat-PR (rabbit polyclonal, Proteintech, 23979-1-AP). Following primary antibody incubation, blots were washed with TBST and incubated with an appropriate HRP secondary antibody (GE Healthcare) for 1 h at room temperature. Signal was detected using Clarity Max Western ECL Substrate (Bio-Rad) and imaged and quantified using Licor Odissey FC/Image Studio software.

### RNA expression analysis

Total RNA was extracted using GenElute total mammalian RNA kit (Sigma) in accordance with the manufacturer’s instructions. First-strand cDNA synthesis was performed using 500 ng of RNA with random primers (ThermoFisher) and MMLV reverse transcriptase (Promega). qRT-PCR was performed using qPCRBIO SyGreen Lo-ROX (PCRbio), and GAPDH was used for normalisation. Primer sequences are provided in a previous study (Shelkovnikova *et al*., 2017).

### In vitro analysis of TDP-43 condensation

*In vitro* TDP-43 clustering analysis with immunodetection and imaging were performed as described in Hodgson, Huang et al., 2024 (doi: 10.2139/ssrn.4721338). Briefly, 1 µM total recombinant TDP-43 (R&D Biosystems, AP-190-100) or supernatant (after centrifuging at 13,300 rpm for 1 min) was mixed with 1 µM poly-PR/GR peptides (as above, BioSynth) or a generic peptide (V5: GKPIPNPLLGLDST, Proteintech) in the assay buffer, and incubated for 10 min. Samples were sedimented and fixed with glutaraldehyde on coverslips, blocked with 1% BSA in PBS for 1 h at RT and incubated with an anti-TDP-43 antibody (1:5000, mouse monoclonal, R&D Biosystems, MAB77781) in the blocking buffer for 2 h. After washes, TDP-43 protein clusters were visualised using anti-mouse AlexaFluor488 antibody (1:2000, ThermoFisher), incubated for 1 h at RT. Coverslips were mounted on a glass slide using Immu-mount (ThermoFisher). Images were taken using Olympus BX57 upright microscope and ORCA-Flash 4.0 camera and processed using cellSens Dimension software (Olympus). Quantification of assemblies was done using the ‘Analyze particles’ tool of Image J. For confirmatory western blot analysis, recombinant TDP-43 samples incubated with or without the peptides as above (for 10 min, and an additional set – for 4 h) were centrifuged at 1,000xg for 10 min. Pellet and supernatant were analysed by western blot using a C-terminal TDP-43 antibody (Sigma).

### Statistical analysis

Analysis was done using respective tests on GraphPad Prism 9 software. N corresponds to the number of biological replicates. Error bars represent S.D. unless indicated otherwise.

## Author contributions

**Rachel E. Hodgson:** Conceptualization; Data curation; Formal analysis; Investigation; Methodology; Writing - original draft; Writing - review and editing. **Jessica Rayment:** Formal analysis; Investigation; Methodology. **Wan-Ping Huang**: Formal analysis; Investigation; Methodology; Writing - review and editing. **Anna Sanchez Avila**: Data curation; Investigation. **Tatyana A. Shelkovnikova:** Conceptualization; Supervision; Funding acquisition; Project administration; Writing - review and editing.

## Disclosure and competing interests statement

The authors declare no competing interests.

## Acknowledgements

This work was supported by the UKRI Future Leaders Fellowship (MR/W004615/1), MRC (MR/W028522/1) and BBSRC (BB/V014110/1) standard grants, and MND Association fellowship/grant (Shelkovnikova/Oct17/968-799). We also acknowledge the MRC grant MR/X012077/1 for Airyscan 2 LMF.

## Notes

### Competing Interest Statement

The authors have declared no competing interest.

### Summary of Updates

Provide full name of authors Amend title and abstract

