## Supplementary material for "Opto-controlled C9orf72 poly-PR forms anisotropic condensates causative of TDP-43 pathology in the nucleus"

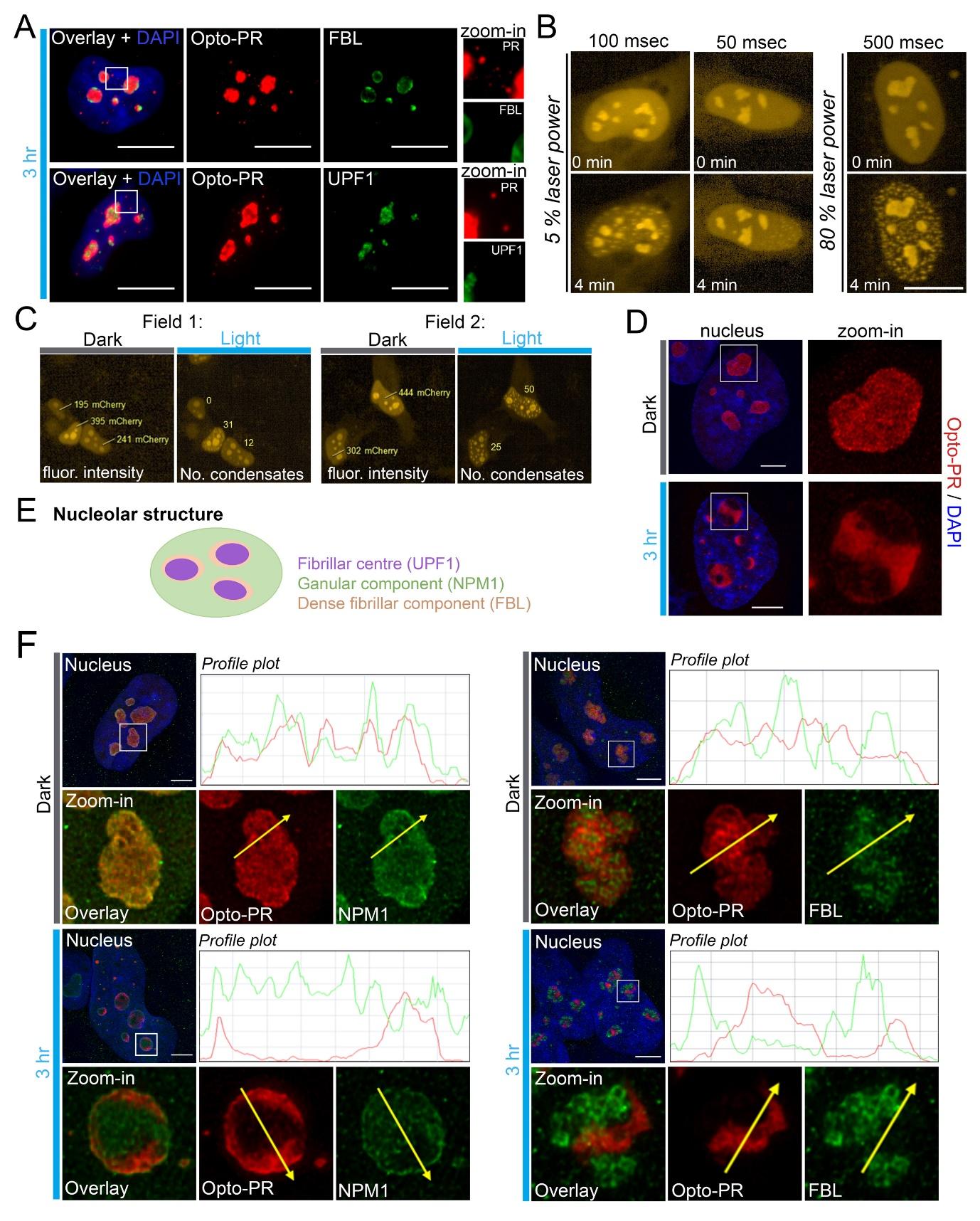

**Figure S1. Validation and characterisation of the opto-PR cellular system.**

(A) Opto-PR condensates are negative for the core nucleolar markers FBL and UPF1. HeLa cells expressing opto-PR were subjected to a 3-h stimulation on the blue-light array. Representative images are shown. Scale bar, 15 µm.

(B) Optimisation of opto-PR condensate induction conditions on a confocal high-content imaging system (Opera Phenix). 488 nm laser was used for stimulation. Representative images are shown. Scale bar, 15 µm.

(C) Assaying the concentration-dependency of opto-PR condensation using individual cell tracking on Opera Phenix. Examples of cells used for correlation analysis are shown. Related to Fig.1G.

(D) Continuous blue-light stimulation induces opto-PR spatial segregation within the nucleolus. Representative images are shown. Scale bar, 5 µm.

(E,F) Opto-stimulation causes opto-PR demixing from NPM1 within the nucleolar granular component (GC), but not from FBL in the dense fibrillar component (DNC). Schematic of the nucleolar organisation (E) and representative images with profile plots (F) are shown. Scale bars, 2 µm.

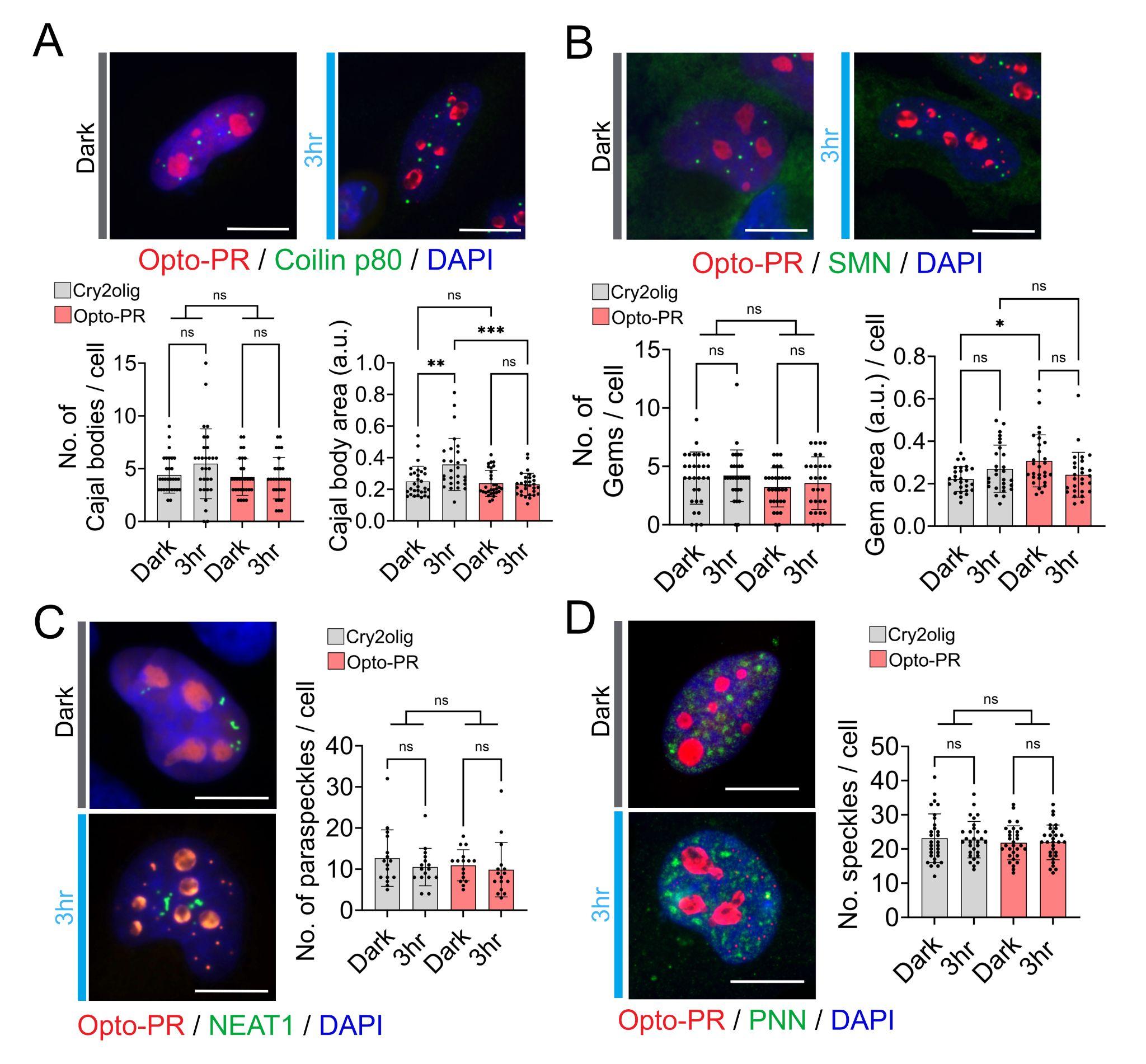

**Figure S2. Effect of diffuse and condensed opto-PR on MLOs.**

(A,B) Opto-PR has a mild effect on Cajal bodies (visualised using coilin p80 as a marker) (A) and Gems (visualised using SMN as a marker) (B) in HeLa cells when diffuse or upon its condensation.

(C,D) Opto-PR does not affect paraspeckles (visualised using NEAT1 lncRNA as a marker) (C) or speckles (visualised using PNN as a marker) (D) in HeLa cells when diffuse or upon its condensation.

Continuous 3-h blue-light stimulation on the blue-light array was used. Representative images and quantification are shown in all panels. Quantification was performed on at least 30 transfected cells per condition from a representative experiment; *p<0.05, **p<0.01, ***p<0.001; one-way ANOVA with Tukey post hoc test. Scale bars, 10 µm.

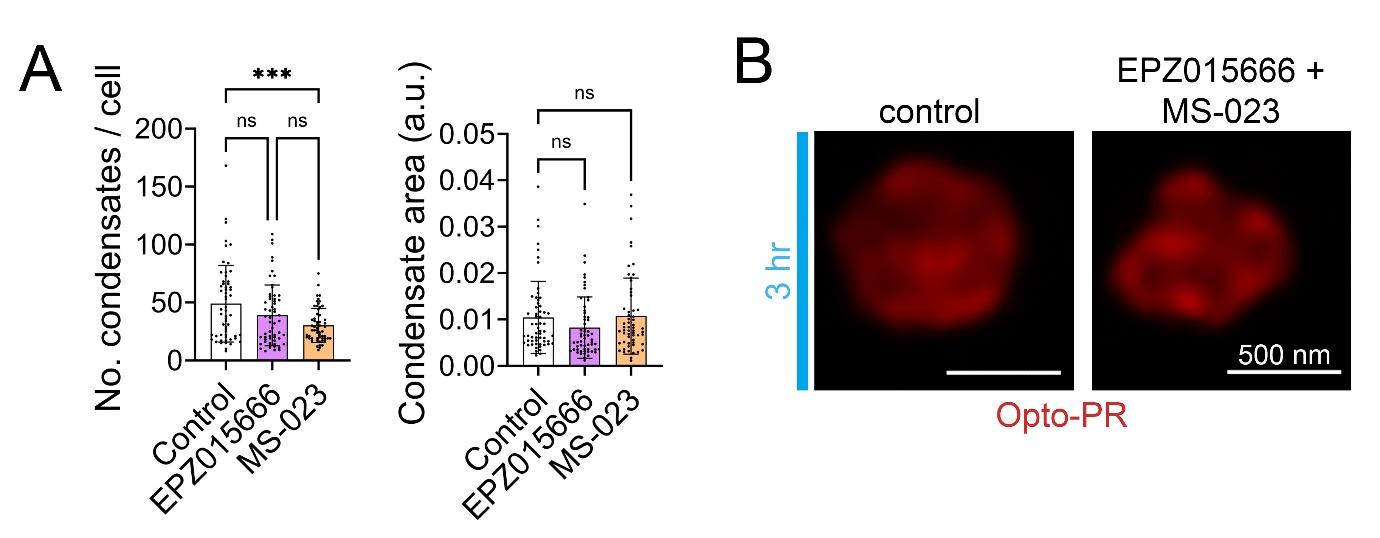

**Figure S3. Arginine dimethylation (DMA) modulates poly-PR condensation in cells.**

(A) Removal of asymmetric arginine dimethylation (aDMA) with MS-023 small molecule inhibitor reduces opto-DPR condensate nucleation. Opto-DPR expressing cells were pre-treated with methyltransferase inhibitors that remove asymmetric (MS023) and symmetric DMA (EPZ015666) for 24 h and subsequently were stimulated with blue light for 3 h continuously. 60 cells were analysed from a representative experiment. ***p<0.001, one-way ANOVA with Dunnett’s post hoc test.

(B) SRM demonstrates unaltered structure of opto-PR condensates in cells pretreated with methyltransferase inhibitors and stimulated as described in A.

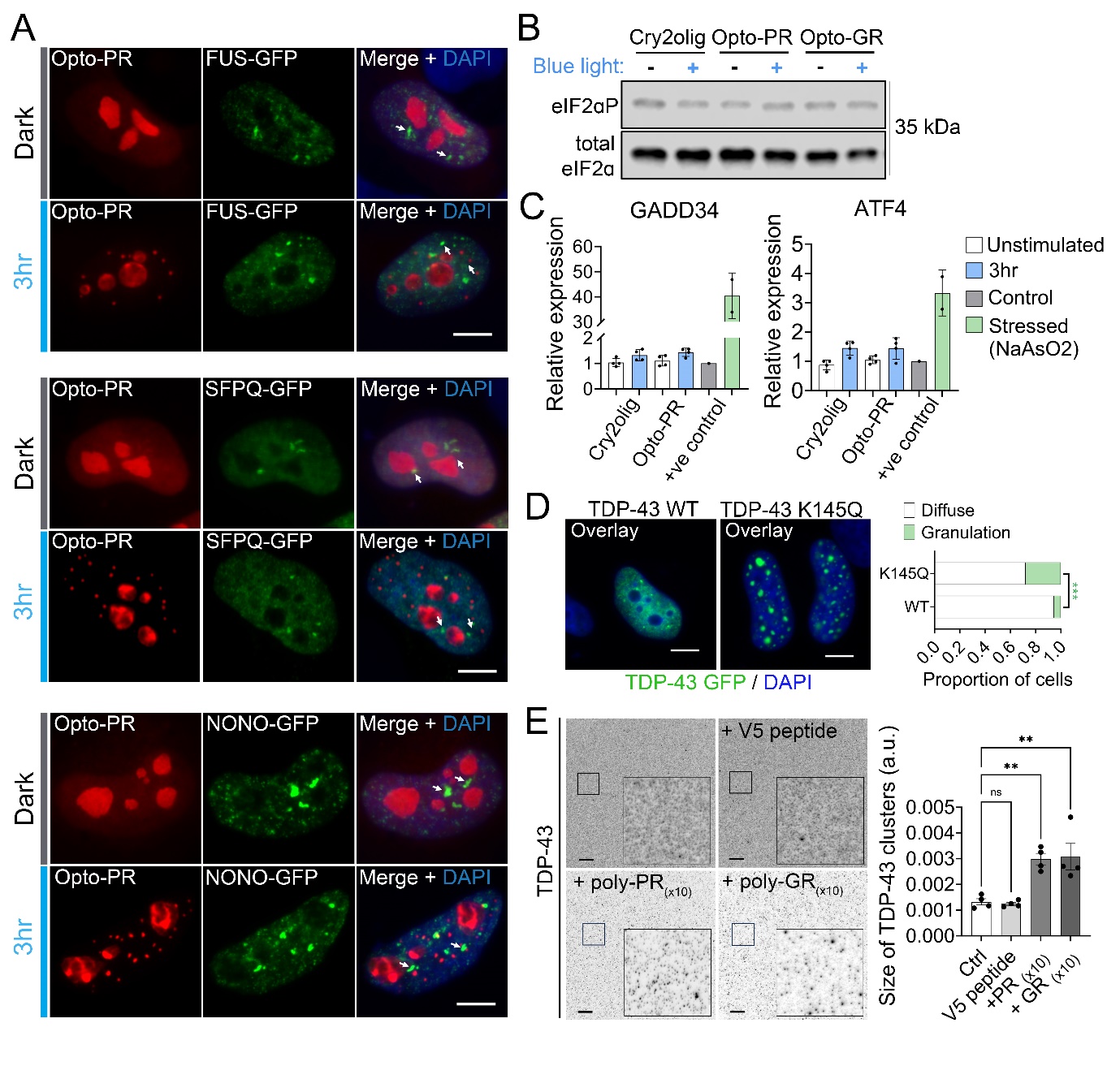

**Figure S4. Effect of opto-PR condensation on RBPs and cellular stress response.**

(A) Opto-PR condensation does not affect the subnuclear distribution of GFP-tagged RBPs FUS, NONO and SFPQ. Note that GFP-positive bright nuclear foci observed both in control and stimulated conditions (arrows) correspond to paraspeckles. Scale bars, 5 µm.

(B) eIF2ɑ phosphorylation at Ser51 is not altered upon opto-PR condensation. Representative western blot of an experiment repeated 3 times is shown.

(C) Opto-PR condensation does not induce upregulation of integrated stress response (ISR) markers GADD34 and ATF4, as analysed by qRT-PCR. Cells stressed with NaAsO_2_ for 1 h and allowed to recover for 3 h were used as a positive control. N=3.

(D) Acetylation-mimic mutation enhances nuclear TDP-43 granulation. Representative images and quantification are shown. >70 cells per condition were analysed from a representative experiment. ***p<0.001, Student’s *t* test. Scale bar, 5 µm.

(E) A generic peptide does not induce TDP-43 clustering *in vitro*. Recombinant TDP-43 (1 µM) was incubated with an equimolar amount of V5 peptide (14-mer, GKPIPNPLLGLDST) and TDP-43 clustering was analysed using fixation/immunostaining as in Fig.3H,I. **p<0.01, ****p<0.0001, one-way ANOVA with Dunnett’s post-hoc test. Scale bar, 10 µm.

In all panels, cells expressing respective constructs for 24 h were subjected to 3-h continuous blue-light stimulation.

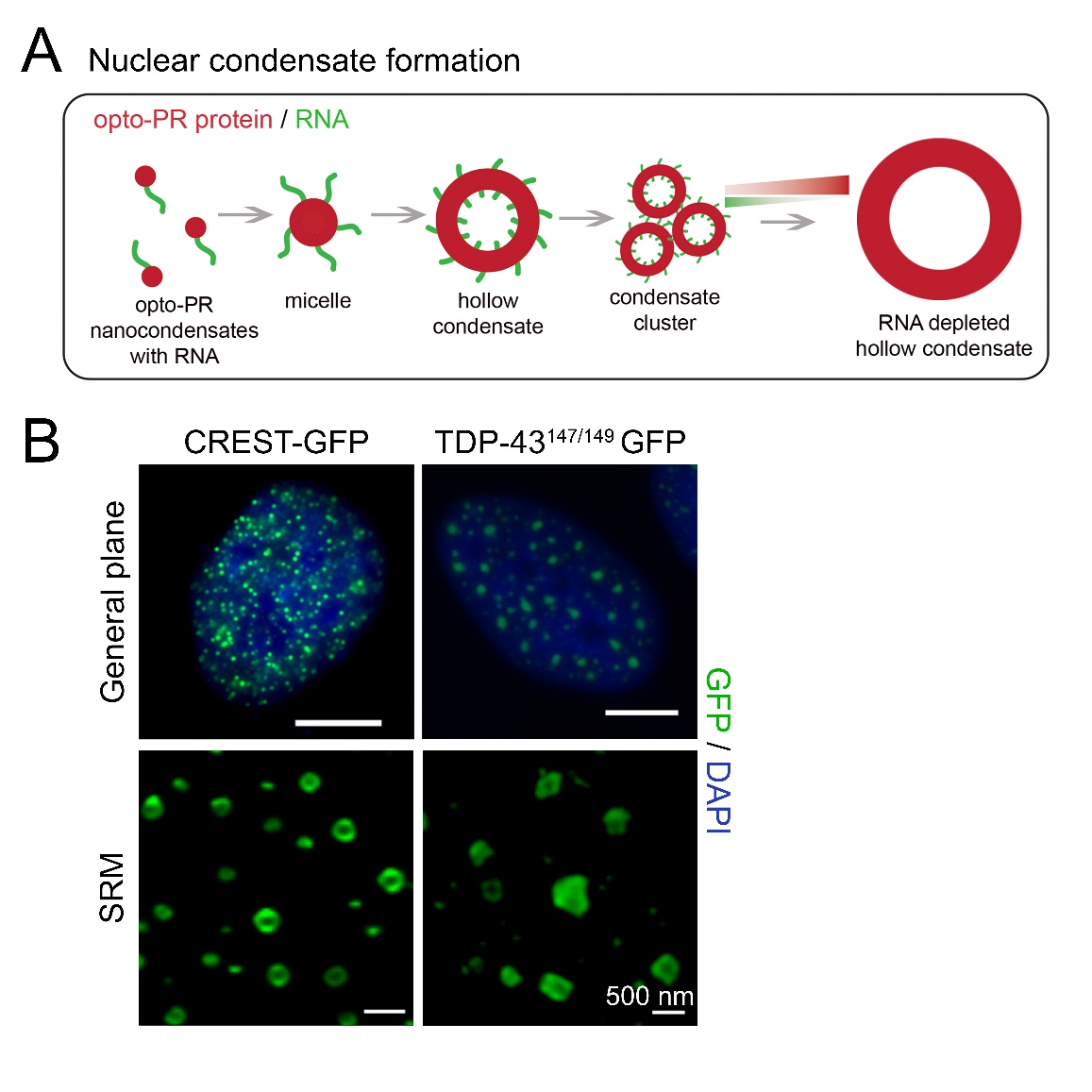

**Figure S5. Anisotropic condensate formation by ALS-linked proteins.**

(A) Model for nuclear opto-PR condensate formation based on (Alshareedah et al., 2020).

(B) Hollow condensate assembly in the nucleus by CREST protein and an RNA binding deficient TDP-43 mutant (F147/147L). Representative images of the cell general plane (conventional microscopy) and close-up (SRM) are shown. Scale bars, 5 µm.
